# High-dimensional Bayesian network inference from systems genetics data using genetic node ordering

**DOI:** 10.1101/501460

**Authors:** Lingfei Wang, Pieter Audenaert, Tom Michoel

## Abstract

Studying the impact of genetic variation on gene regulatory networks is essential to understand the biological mechanisms by which genetic variation causes variation in phenotypes. Bayesian networks provide an elegant statistical approach for multi-trait genetic mapping and modelling causal trait relationships. However, inferring Bayesian gene networks from high-dimensional genetics and genomics data is challenging, because the number of possible networks scales super-exponentially with the number of nodes, and the computational cost of conventional Bayesian network inference methods quickly becomes prohibitive. We propose an alternative method to infer high-quality Bayesian gene networks that easily scales to thousands of genes. Our method first reconstructs a node ordering by conducting pairwise causal inference tests between genes, which then allows to infer a Bayesian network via a series of independent variable selection problems, one for each gene. We demonstrate using simulated and real systems genetics data that this results in a Bayesian network with equal, and sometimes better, likelihood than the conventional methods, while having a significantly higher overlap with groundtruth networks and being orders of magnitude faster. Moreover our method allows for a unified false discovery rate control across genes and individual edges, and thus a rigorous and easily interpretable way for tuning the sparsity level of the inferred network. Bayesian network inference using pairwise node ordering is a highly efficient approach for reconstructing gene regulatory networks when prior information for the inclusion of edges exists or can be inferred from the available data.

## 1 Introduction

Complex traits and diseases are driven by large numbers of genetic variants, mainly located in non-coding, regulatory DNA regions, affecting the status of gene regulatory networks [1–5]. While important progress has been made in the experimental mapping of protein-protein and protein-DNA interactions [6–8], the context-specific and dynamic nature of these interactions means that comprehensive, experimentally validated, cell-type or tissue-specific gene networks are not readily available for human or animal model systems. Furthermore, knowledge of physical protein-DNA interactions does not always allow to predict functional effects on target gene expression [9]. Hence, statistical and computational methods are essential to reconstruct context-specific, causal, trait-associated networks by integrating genotype and gene, protein and/or metabolite expression data from a large number of individuals segregating for the traits of interest [1–3].

Gene network inference is a deeply studied problem in computational biology [10–17]. Among the many successful methods that have been devised, Bayesian networks are a particularly powerful approach for modelling causal relationships and incorporating prior knowledge [10, 18–22]. In the context of complex trait genetics, Bayesian networks are a natural generalization of linear models for mapping the genetic architecture of a single traits to the modelling of conditional independence and causal dependence between multiple traits, including molecular abundance traits [23–27]. They have therefore become the most popular method for modelling joint genetic linkages and gene networks using systems genetics data. Among their many successes, Bayesian networks have been used to identify key driver genes of type 1 diabetes [28], Alzheimer’s disease [29, 30], temporal lobe epilepsy [31] and cardiovascular disease [32]. However, Bayesian network inference is computationally demanding and limited to relatively small-scale systems. In this paper we address the question whether Bayesian network inference from genotype and gene expression data is feasible on a truely transcriptome-wide scale without sacrificing performance in terms of model fit and overlap with known interactions.

A Bayesian gene network consists of a directed graph without cycles, which connects regulatory genes to their targets, and which encodes conditional independence between genes. The structure of a Bayesian network is usually inferred from the data using score-based or constraint-based approaches [21]. Score-based approaches maximize the likelihood of the model, or sample from the posterior distribution using Markov chain Monte Carlo (MCMC), using edge additions, deletions or inversions to search the space of network structures. Score-based methods have been shown to perform well using simulated genetics and genomics data [33, 34]. Constraint-based approaches first learn the undirected skeleton of the network using repeated conditional independence tests, and then assign edge directions by resolving directional constraints (v-structures and acyclicity) on the skeleton. They have been used for instance in the joint genetic mapping of multiple complex traits [27]. However, the computational cost of both approaches is high. Because the number of possible graphs scales super-exponentially with the number of nodes, Bayesian gene network inference with conventional methods is feasible for systems of at most a few hundred genes or traits, and usually requires a hard limit on the number of regulators a gene can have as well as a preliminary dimension reduction step, such as filtering or clustering genes based on their expression profiles [24, 29, 30, 32].

Modern sequencing technologies however generate transcript abundance data for ten-thousands of coding and non-coding genes, and large sample sizes mean that ever more of those are detected as variable across individuals [35–37]. Moreover, to explain why genetic associations are spread across most of the genome, a recently proposed “omnigenic” model of complex traits posits that gene regulatory networks are sufficiently interconnected such that all genes expressed in a disease or trait-relevant cell or tissue type affect the functions of core trait-related genes [5]. The limitations of current Bayesian gene network inference methods mean that this model can be neither tested nor accomodated. Hence there is a clear and unmet need to infer Bayesian networks from very high-dimensional systems genetics data.

Here we propose a novel method to infer high-quality causal gene networks that scales easily to ten-thousands of genes. Our method is based on the fact that if an ordering of nodes is given, such that the parents of any node must be a subset of the predecessors of that node in the given ordering, then Bayesian network inference reduces to a series of independent variable or feature selection problems, one for each node [21, 38]. While reconstructing a node ordering is challenging in most application domains, *pairwise* comparisons between nodes can sometimes be obtained. If prior information is available for the likely inclusion of every edge, our method ranks edges according to the strength of their prior evidence (e.g. p-value) and incrementally assembles them in a directed acyclic graph, which defines a node ordering, by skipping edges that would introduce a cycle. Prior pairwise knowledge in systems biology includes the existence of TF binding motifs [39], or known protein-DNA and protein-protein interactions [40, 41], and those have been used together with score-based MCMC methods in Bayesian network inference previously [19, 20].

In systems genetics, where genotype and gene expression data are available for the same samples, instead of using external prior interaction data, pairwise causal inference methods can be used to estimate the likelihood of a causal interaction between every pair of genes [42–48]. To accomodate the fact that the same gene expression data is used to derive the node ordering and subsequent Bayesian network inference, we propose a novel generative model for genotype and gene expression data, given the structure of a gene regulatory graph, whose log-likelihood decomposes as a sum of the standard log-likelihood for observing the expression data and a term involving the pairwise causal inference results. Our method can then be interpreted as a greedy optimization of the posterior log-likelihood of this generative model.

## 2 Methods

### 2.1 An algorithm for the inference of gene regulatory networks from systems genetics data

To allow the inference of gene regulatory networks from high-dimensional systems genetics data, we developed a method that exploits recent algorithmic developments for highly efficient mapping of expression quantitative trait loci (eQTL) and pairwise causal interactions. A general overview of the method is given here, with concrete procedures for every step detailed in subsequent sections below.

#### A. eQTL mapping

When genome-wide genotype and gene expression data are sampled from the same unrelated individuals, fast matrix-multiplication based methods allow for the efficient identification of statistically significant eQTL associations [49–52]. Our method takes as input a list of genes, and for every gene its most strongly associated eQTL (Figure 1A). Typically only *cis*-acting eQTLs (i.e. genetic variants located near the gene of interest) are considered for this step, but this is not a formal requirement. Multiple genes can have the same associated eQTL, and genes without significant eQTL can be included as well, although these will only be allowed to have incoming edges in the resultant Bayesian networks.

**Figure 1:**
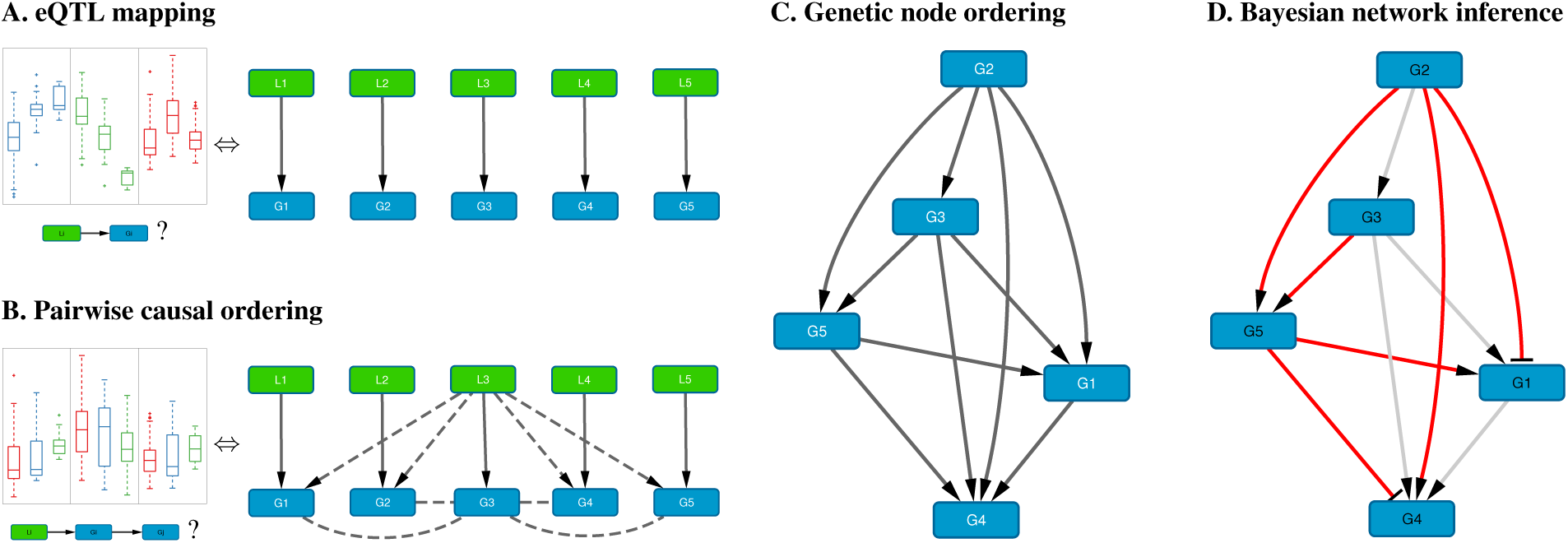
Schematic overview of the method. **A.** For each gene *G*_*i*_, the *cis*-eQTL *L*_*i*_ whose genotype explains most of the variation in *G*_*i*_ expression is calculated; shown on the left are typical eQTL associations for three genes (colored blue, green and red) where each box shows the distribution of expression values for samples having a particular genotype for that gene’s eQTL. **B.** Pairwise causal inference is carried out which considers in turn each gene *G*_*i*_ and its eQTL *L*_*i*_ to calculate the likelihood of this gene being causal for all others; shown on the left is a typical example where an eQTL *L*_*i*_ is associated with expression of *G*_*i*_ (red) and with expression of a correlated gene *G*_*j*_ (blue), but not with expression of *G*_*j*_ adjusted for *G*_*i*_ (green), resulting in a high likelihood score for the causal ordering *G*_*i*_ →*G*_*j*_. **C.** A maximum-weight directed acyclic graph having the genes as its nodes is derived from the pairwise causal interactions, which induces a “genetic” node ordering. **D.** Variable selection is used to determine a sparse Bayesian gene network, which must be a sub-graph of the maximum-weight graph (red edges, Bayesian network; gray edges, causal orderings deemed not significant or indirect by the variable selection procedure); the signs of the maximum-likelihood linear regression coefficients determine whether an edge is activating (arrows) or repressing (blunt tips).

#### B. Pairwise causal ordering

Given a set of genes and their respective eQTLs, pairwise causal interactions between all genes are inferred using the eQTLs as instrumental variables (Figure 1B). While there is a great amount of literature on this subject (cf. Introduction), only two stand-alone software packages are readily available: CIT [47] and Findr [48]. In our experience, only Findr is sufficiently efficient to test for causality between millions of gene pairs.

#### C. Genetic node ordering

In Section 2.3 we introduce a generative probabilistic model for jointly observing eQTL genotypes and gene expression levels given the structure of a gene regulatory network. In this model, the posterior log-likelihood of the network given the data decomposes as a sum of two terms, one measuring the fit of the undirected network to the correlation structure of the gene ex-pression data, and the other measuring the fit of the edge directions to the pairwise causal interactions inferred using the eQTLs as instrumental variables. The latter is optimized by a maximum-weight directed acyclic graph (DAG), which induces a topological node ordering, which we term “genetic node ordering” in reference to the use of individual-level genotype data to orient pairs of gene expression traits (Figure 1C).

#### D. Bayesian network inference

The genetic node ordering fixes the directions of the Bayesian network edges. Variable selection methods are then used to determine the optimal sparse representation of the inverse covariance matrix of the gene expression data by a subgraph of the maximum-weight DAG (Figure 1D). In this paper, we consider two approaches: *(i)* a truncation of the pairwise interaction scores retaining only the most confident (highest weight) edges in the maximum-weight DAG, and *(ii)* a multi-variate, L1-penalized lasso regression [53] to select upstream regulators for every gene. Given a sparse DAG, maximum-likelihood linear regression is used to determine the input functions and whether an edge is activating or repressing.

### 2.2 Bayesian network model with prior edge information

A Bayesian network with *n* nodes (random variables) is defined by a DAG *𝒢* such that the joint distribution of the variables decomposes as

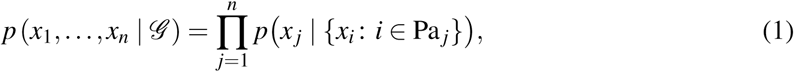

where Pa _*j*_ denotes the set of parent nodes of node *j* in the graph *𝒢*. We only consider linear Gaussian networks [21], where the conditional distributions are given by normal distributions whose means depend linearly on the parent values (see Supplementary Information).

The likelihood of observing a data matrix **X** ∈ R^*n*×*m*^ with expression levels of *n* genes in *m* independent samples given a DAG *𝒢* is computed as

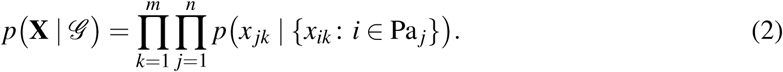

Using Bayes’ theorem we can then write the likelihood of observing *𝒢* given the data **X**, upto a normalization constant, as

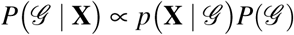

where *P*(*𝒢*) is the prior probability of observing *𝒢*. Note that we use a lower-case ‘*p*’ to denote probability density functions and upper-case ‘*P*’ to denote discrete probability distributions.

Our method is applicable if pairwise prior information is available, i.e. for prior distributions satisfying

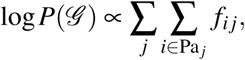

with *f*_*ij*_ a set of non-negative weights that are monotonously increasing in our prior belief that there exists a directed edge from node *i* to node *j* (e.g. *f*_*ij*_ ∝‒log *p*_*ij*_, where *p*_*ij*_ is a *p*-value). Note that setting *f*_*ij*_ = 0 excludes the edge (*i, j*) from being present in *𝒢*.

### 2.3 Bayesian network model for systems genetics data

When genotype and gene expression data are available for the same samples, instrumental variable methods can be used to infer the likelihood of a causal interaction between every pair of genes [42–48]. Previously, such pairwise probabilities have been used as priors in conventional score-based Bayesian network inference [23, 33], but this is unsatisfactory, because a prior, by definition, s hould n ot be inferred from the same expression data that is used to learn the model. Other methods have addressed this by augmenting the gene network model with genotypic variables [25, 26], but this increases the size and complexity of the model even further. Here we introduce a model to use pairwise causal inference that does not suffer from these limitations.

Let *𝒢* and **X** again be a DAG and a matrix of gene expression data for *n* genes, respectively, and let **E** ∈R^*n*×*m*^ be a matrix of genotype data for the same samples. For simplicity we assume that each gene has one associated genotypic variable (e.g. its most significant *cis*-eQTL), but this can be extended easily to having more than one eQTL per gene or to some genes having no eQTLs. Using the rules of conditional probability, the joint probability (density) of observing **X** and **E** given *𝒢* can be written, upto a normalization constant, as

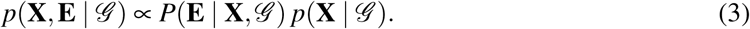

The distribution *p*(**X** | *𝒢*) is obtained from the standard Bayesian network equations [eq. (2)], and we define the conditional probability of observing **E** given **X** and *𝒢* as

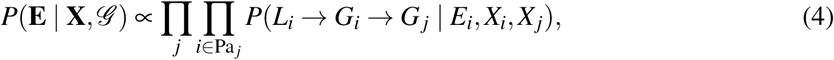

where *E*_*i*_, *X*_*i*_ ∈ R^*m*^ are the *i*th rows of **E** and **X**, respectively, and *P*(*L*_*i*_ → *G*_*i*_→ *G*_*j*_ | *E*_*i*_, *X*_*i*_, *X*_*j*_) is the probability of a causal interaction from gene *G*_*i*_ to *G*_*j*_ inferred using *G*_*i*_’s eQTL *L*_*i*_ as a causal anchor. In other words, conditional on a gene-to-gene DAG *𝒢* and a gene expression data matrix, our model assumes that it is more likely to observe genotype data that would lead to causal inferences consistent with *𝒢* than data that would lead to inconsistent inferences. Other variations on this model can be considered as well, for instance one can include a penalty for interactions that are not present in the graph, as long as the final model can be expressed in the form

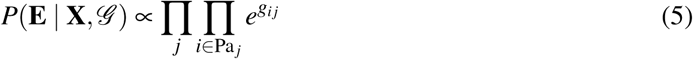

with *g*_*ij*_ monotonously increasing in the likelihood of a causal inference *L*_*i*_ →*G*_*i*_ →*G*_*j*_.

Combining eqs. (3) and (5) with Bayes’ theorem and a uniform prior *P*(*𝒢*) = const, leads to an expression of the posterior log-likelihood that is formally identical to the model with prior edge information,

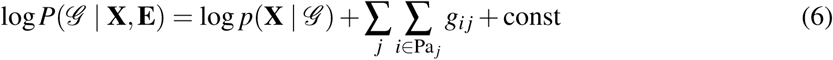

As before, if *g*_*ij*_ = 0, the edge (*i,j*) is excluded from being part of *𝒢*; this would happen for instance if gene *i* has no associated genotypic variables and consequently zero probability of being causal for any other genes given the available data. Naturally, informative pairwise graph priors of the form 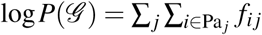, can still be added to the model, when such information is available.

### 2.4 Bayesian network parameter inference

Given a DAG *𝒢*, the maximum-likelihood parameters of the conditional distributions [eq. (1)], in the case of linear Gaussian networks, are obtained by linear regression of a gene on its parents’ expression profiles (see Supplementary Information). For a specific DAG, we will use the term “Bayesian network” to refer to both the DAG itself as well as the probability distribution induced by the DAG with its maximum-likelihood parameters.

### 2.5 Reconstruction of the node ordering

Without further sparsity constraints in eq. (6), and again assuming for simplicity that each gene has exactly one eQTL, the log-likelihoodims aximizedby a DAG with *n*(*n* ‒ 1)*/*2 edges. Such a DAG *𝒢* defines a node ordering ≺ where *i* ≺ *j* ⇔ *i* ∈ Pa*j*. Standard results in Bayesian network theory show that for a linear Gaussian network, the likelihood function (2) is invariant under arbitrary changes of the node ordering (see [21] and Supplementary Information). Hence to maximize eq. (6) we need to find the node ordering or DAG which maximizes the term 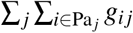. Finding the maximum-weight DAG is an NP-hard problem with no known polynomial approximation algorithms with a strong guaranteed error bound [54, 55]. We therefore employed a greedy algorithm, where given *n* genes and the log-likelihood *g*_*i j*_ of regulation between every pair of them, we first rank the regulations according to their likelihood. The regulations are then added to an empty network one at a time starting from the most probable one, but avoiding those that would create a cycle, until a maximum-weight DAG with *n*(*n*‒1)*/*2 edges is obtained. Other edges are assigned probability 0 to indicate exclusion. The heuristic maximum-weight DAG reconstruction was implemented in Findr [48] as the command netr one greedy, with the *vertex-guided* algorithm for cycle detection [56].

### 2.6 Causal inference of pairwise gene regulations

We used Findr 1.0.6 (pij gassist function) [48] to perform causal inference of gene regulatory interactions based on gene expression and genotype variation data. For every gene, its strongest *cis*-eQTL was used as a causal anchor to infer the probability of regulation between that gene and every other gene. Findr outputs posterior probabilities *P*_*ij*_ (i.e. one minus local FDR), which served directly as weights in model (6), i.e. we set *g*_*ij*_ = log *P*_*ij*_. To verify the contribution from the inferred pairwise regulations, we also generated random pairwise probability matrices which were treated in the same way as the informative ones in the downstream analyses.

### 2.7 Findr and random Bayesian networks from node orderings

The node ordering reconstruction removes less probable, cyclic edges, and results in a (heuristic) maximum-weight DAG *𝒢* with edge weights 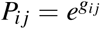. We term these weighted DAGs as *findr* or *random Bayesian networks*, depending on the pairwise information used. A significance threshold can be applied on the continuous networks, to convert them to binary Bayesian networks at any desired sparsity level and thereby perform variable selection for the parents of every gene.

### 2.8 Lasso-findr and lasso-random Bayesian networks using penalized regression on ordered nodes

As a second approach to perform variable selection in the maximum-weight DAGs, we performed hypothesis testing for every gene on whether each of its predecessors (in the findr or random Bayesian network) is a regulator, using L1-penalized lasso regression [53] with the lassopv package [57] (see Supplementary Information). We calculated for every regulator the p-value of the critical regularization strength when the regulator first becomes active in the lasso path. This again forms a continuous Bayesian network in which smaller p-values indicate stronger significance. These Bayesian networks were termed the *lasso-findr* and *lasso-random Bayesian networks*.

### 2.9 Score-based bnlearn-hc and constraint-based bnlearn-fi Bayesian networks from package bnlearn

For comparison with score-based Bayesian network inference methods, we applied the hc function of the R package bnlearn [58], using the Akaike information criterion (AIC) penalty to enforce sparsity. This algorithm starts from a random Bayesian network and iteratively performs greedy revisions on the network to reach a local optimum of the penalized likelihood function. Since the log-likelihood is equivalent to minus the average (over nodes) log unexplained variance (see Supplementary Information), which diverges when the number of regulators exceeds the number of samples, we enforced the number of regulators for every gene to be smaller than 80% of the number of samples. For each AIC penalty, one hundred random restarts were carried out and only the network with highest likelihood score was selected for downstream analyses. These Bayesian networks were termed the *bnlearn-hc* Bayesian networks.

For comparison with constraint-based Bayesian network inference methods (e.g. [59]), we applied the fast.iamb function of the R package bnlearn [58], using nominal type I error rate. These Bayesian networks were termed the *bnlearn-fi* Bayesian networks.

To account for the role and information of cis-eQTLs on gene expression, we also included the strongest cis-eQTL of every gene in the bnlearn-based network reconstructions, for an approach similar to [25, 26, 34]. Cis-eQTLs were only allowed to have out-going edges, using the blacklist function in bnlearn. We then removed cis-eQTL nodes from the reconstructed networks, resulting in Bayesian gene networks termed *bnlearn-hc-g* and *bnlearn-fi-g* respectively.

### 2.10 Evaluation of false discovery control in network inference

An inconsistent false discovery control (FDC) reduces the overall accuracy of the reconstructed network [57]. We empirically evaluated the FDC using a linearity test on genes that are both targets and regulators. The linearity test assesses whether the number of false positive regulators for each gene increases linearly with the number of candidate regulators, a consequence of consistent FDC. The top 5% predictions were discarded to remove genuine interactions. See [57] for method details.

### 2.11 Precision-recall curves and points

We compared reconstructed Bayesian networks with gold standards using precision-recall curves and points, for continuous and binary networks respectively. For Geuvadis datasets, we only included regulator and target genes that are present in both the transcriptomic dataset and the gold standard.

### 2.12 Assessment of predictive power for Bayesian networks

To assess the predictive power of different Bayesian network inference methods, we used five-fold cross-validation to compute the training and testing errors from each method, in terms of the root mean squared error (rmse) and mean log squared error (mlse) across all genes in all testing data (Supplementary Information, Algorithm S1). For continuous Bayesian networks from non-bnlearn methods, we applied different significance thresholds to obtain multiple binary Bayesian networks that form a curve of prediction errors.

### 2.13 Data

We used the following datasets to infer and evaluate Bayesian gene networks:

- The DREAM 5 Systems Genetics challenge A (DREAM) provided a unique testbed for network inference methods that utilize genetic variations in a population (https://www.synapse.org/\#!Synapse:syn2820440/wiki/). The DREAM challenge included 15 simulated datasets of expression levels of 1000 genes and their best eQTL variations. To match the high-dimensional property of real datasets where the number of genes exceeds the number of individuals, we analyzed datasets 1, 3, and 5 with 100 individuals each. Around 25% of the genes within each dataset had a cis-eQTL, defined in DREAM as directly affecting the expression level of the corresponding gene. Since the identity of cis-eQTLs is not revealed, we used kruX [60] to identify them, allowing for one false discovery per dataset. The DREAM challenge further provides the groundtruth network for each dataset, varying from around 1000 to 5000 interactions.
- The Geuvadis consortium is a population study providing RNA sequencing and genotype data of lymphoblastoid cell lines in 465 individuals. We obtained gene expression levels and genotype information, as well as the eQTL mapping from the original study [61]. We limited our analysis to 360 European individuals, and after quality control, a total of 3172 genes with significant cis-eQTLs remained. To validate the inferred gene regulatory networks from the Geuvadis dataset, we obtained three groundtruth networks: (1) differential expression data from siRNA silencing experiments of transcription-associated factors (TFs) in a lymphoblastoid cell line (GM12878) [62]; (2) DNA-binding information of TFs in the same cell line [62]; (3) the filtered proximal TF-target network from [7]. The Geuvadis dataset overlapped with 6,790 target genes, and 6 siRNA-targeted TFs and 20 DNA-binding TFs in groundtruth 1 and 2, respectively, and with 7,000 target genes and 14 TFs in groundtruth 3.

We preprocessed all expression data by converting them to a standard normal distribution separately for each gene, as explained in [48].

## 3 Results

### 3.1 Genetic node ordering permits high-dimensional Bayesian network inference

We developed a method for Bayesian network inference from high-dimensional systems genetics data which reconstructs a maximum-weight directed acyclic graph (DAG) from the confidence scores of pairwise causal inferences between gene expression traits using eQTLs as causal anchors, and which uses the node ordering induced by this DAG (termed “genetic node ordering” in reference to the use of genotype data to orient network edges) to decompose the Bayesian network inference task into a series of independent variable selection problems (Methods, Section 2.1, Figure 1). Using an efficient implementation for the causal inference step [48], this approach allows to reconstruct Bayesian networks with thousands to ten-thousands of nodes. Our method is based on score-based Bayesian network inference methods for systems with pre-defined node orderings [21,38], but differs in that the ordering is inferred from the same expression data, augmented with matched genotype data from the same samples, that is used for the subsequent Bayesian network log-likelihood maximization, using a single generative model (Methods Section 2.3), rather than relying on external prior information to determine the node ordering. Its computational efficiency is due to restricting the graph structure search space to Bayesian gene networks compatible with this inferred node ordering. This differs substantially from conventional score-based and constraint-based methods, including those that use genotype and gene expression data [25, 26, 34], where the search space can only be reduced by limiting the possible number of parents for each gene to an artificially small number [21]. For clarity, a comparison of the main characteristics of the Bayesian network inference approaches considered in this paper is included in Supplementary Table S1.

### 3.2 Lasso-findr Bayesian networks correctly control false discoveries

We inferred findr and lasso-findr Bayesian networks for the DREAM datasets, using Findr and lassopv respectively (Methods). The Findr method predicts targets for each regulator using a local FDR score [63] which allows false discovery control (FDC) for either the entire regulator-by-target matrix, or for a specific regulator of interest [43, 48]. However, the enforcement of a gene ordering/Bayesian network partly broke the FDC, as seen from the linearity test (Methods) in Figure 2A. By performing an extra lasso regression on top of the acyclic findr network, proper FDC was restored in the lasso-findr Bayesian network (Figure 2B, Supplementary Figure S1).

**Figure 2:**
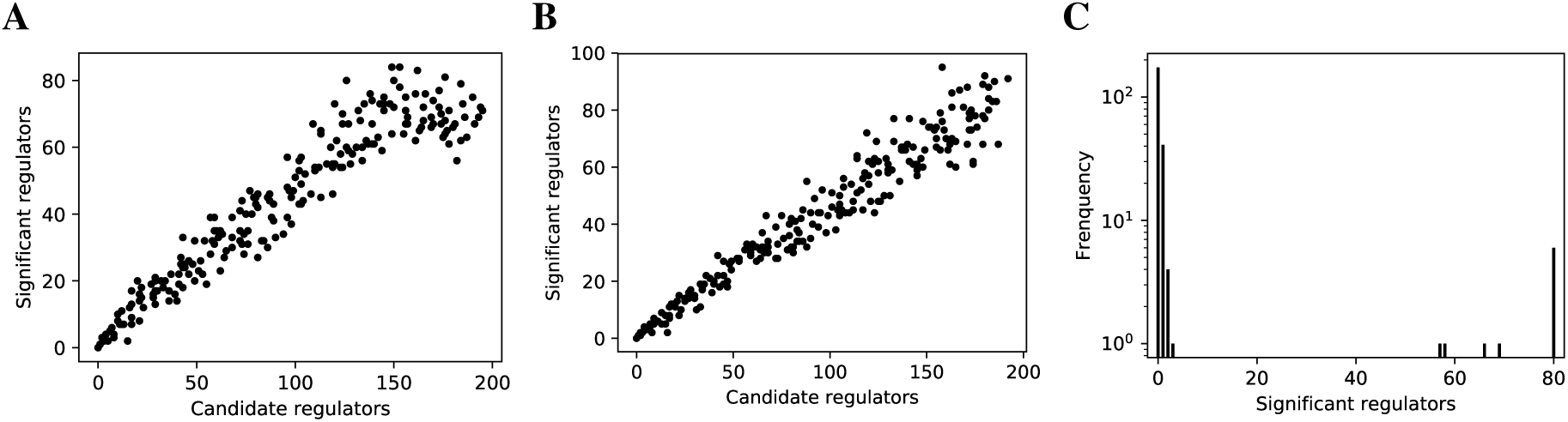
False discovery controls of different Bayesian networks. (**A, B**) The linearity test of findr (**A**) and lasso-findr (**B**) Bayesian networks at 10,000 significant interactions on DREAM dataset 1. (**C**) The histogram of significant regulator counts for each target gene in the bnlearn-hc Bayesian network with AIC penalty 8 on DREAM dataset 1.

In contrast, score-based bnlearn-hc Bayesian networks (Methods), inferred from multiple DREAM datasets and for a spectrum of network sparsities (AIC penalty strengths from 8 to 12 in steps of 0.5), displayed a highly skewed in-degree distribution, with most genes having few regulators, but several with near 80 regulators each, i.e. the maximum allowed (Figure 2C, Supplementary Figure S2). This indicates that score-based Bayesian networks lack a unified FDR control, i.e. that each gene retained incoming interactions at different FDR levels. We believe this is due to the log-likelihood score function employed by bnlearn-hc. Since the log-likelihood corresponds to the average logarithm of the unexplained variance, this score intrinsically tends to focus on the explanation of variances from a few variables/genes, especially in high-dimensional settings where this can lead to arbitrarily large score values (see Supplementary Information). Using the total proportion of explained variance as the score may spread regulations over more target genes, but this score is not implemented in bnlearn.

Constraint-based bnlearn-fi Bayesian networks (Methods) did not allow for unbiased FDC either, as they do not have a fully adjustable sparsity level. We varied its “nominal type I error rate” from 0.001 to 0.2, but the number of significant interactions varied very little on DREAM dataset 1 (Supplementary Figure S3).

Incorporating genotypic information in score-based (bnlearn-hc-g) or constraint-based (bnlearn-fi-g) Bayesian networks did not resolve these issues, as the problems of lacking FDC and oversparsity persisted (Supplementary Figure S4, Supplementary Figure S5).

### 3.3 Findr and lasso Bayesian networks recover genuine interactions more accurately than MCMC or constraint-based networks

We compared the inferred Bayesian networks from all methods against the groundtruth network of the DREAM challenge. We drew precision-recall (PR) curves, or points for the binary Bayesian networks from bnlearn-based methods. As shown in Figure 3, the findr, lasso-findr, and lasso-random Bayesian networks were more accurate predictors of the underlying network structure. The inclusion of genotypic information improved the precision of bnlearn methods, but it remained less optimal than findr and lasso-based Bayesian networks.

**Figure 3:**
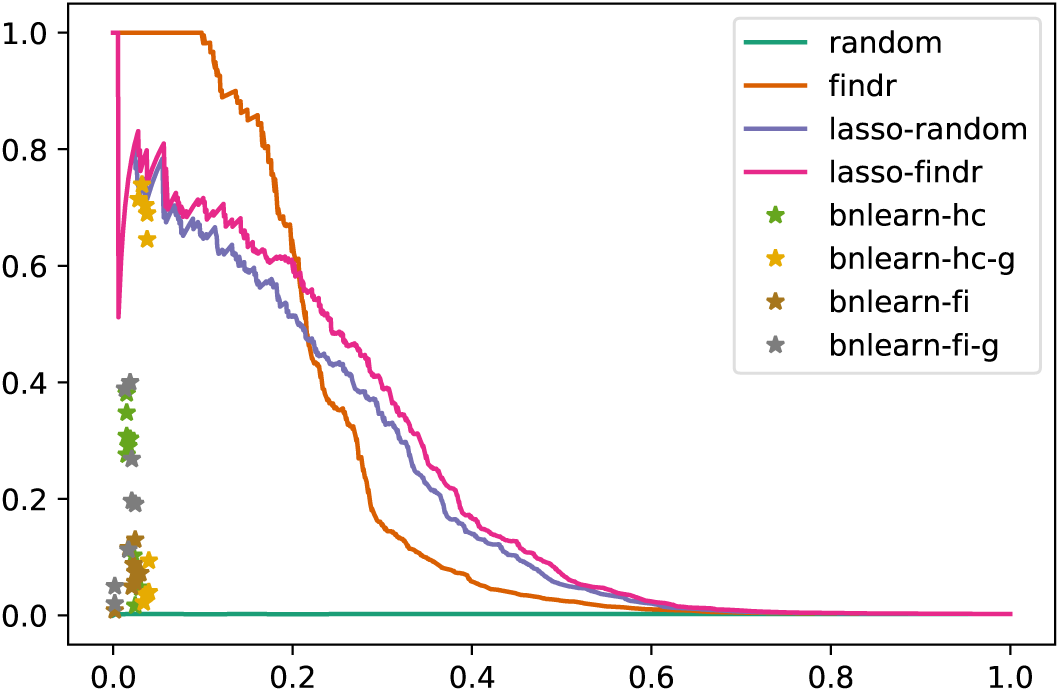
Precision-recall curves/points of reconstructed Bayesian networks for DREAM dataset 1.

### 3.4 Findr and lasso Bayesian networks obtain superior predictive performances

We validated the predictive performances of all networks in the structural equation context (see Supplementary Information). Under 5-fold cross validation, a linear regression model for each gene on its parents is trained based on the Bayesian network structure inferred from each training set, to predict expression levels of all genes in the test set (Methods). Predictive errors were measured in terms of root mean squared error (rmse) and mean log squared error (mlse; the score optimized by bnlearn-hc). The findr Bayesian network explained the highest proportion of expression variation (≈ 2%) in the test data and identified the highest number of regulations (200 to 300), with runners up from lasso-based networks (≈ 1% variation, 50 regulations, Figure 4). The explained variance by findr and lasso networks grew to ≈10% when more samples were added (DREAM dataset 11 with 999 samples, Supplementary Figure S6). Training errors did not show overfitting of predictive performances in the test data (Supplementary Figure S7).

**Figure 4:**
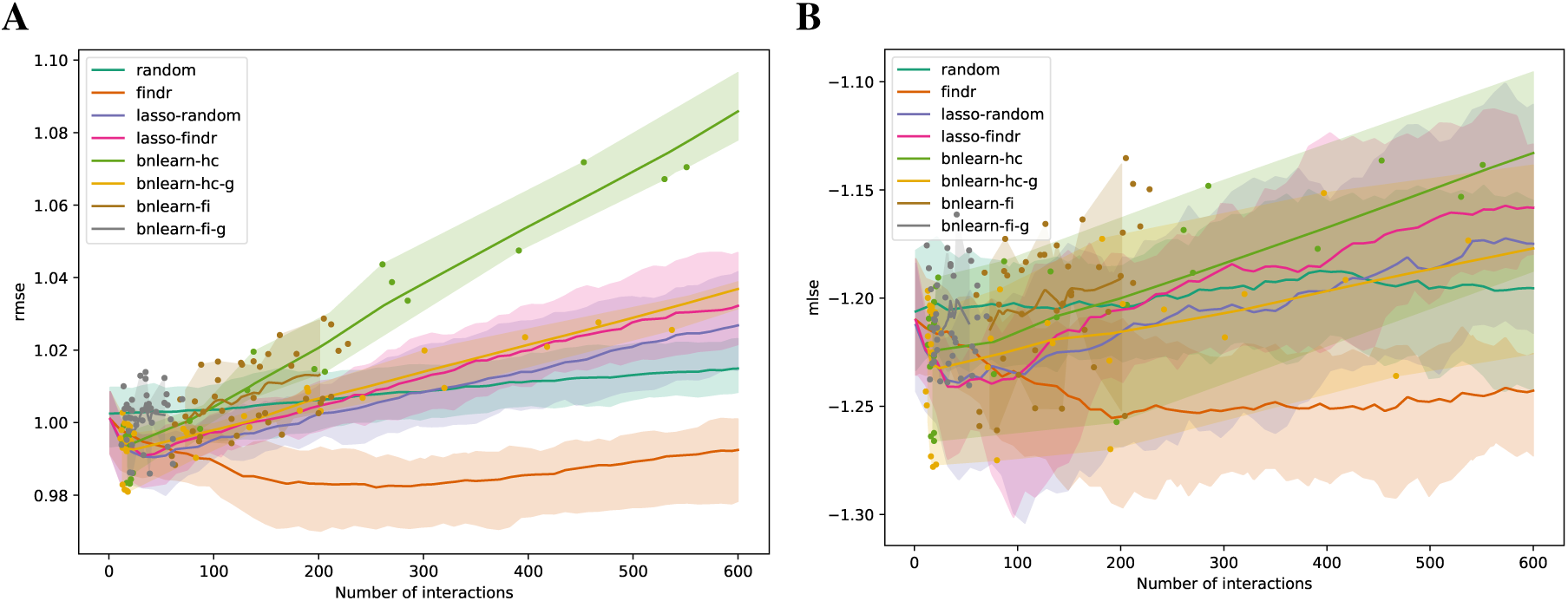
The root mean squared error (rmse, **A**) and mean log squared error (mlse, **B**) in test data are shown as functions of the numbers of predicted interactions in five-fold cross validations using linear regression models. Shades and lines indicate minimum/maximum values and means respectively. Root mean squared errors greater than 1 indicate over-fitting. DREAM dataset 1 with 100 samples was used.

### 3.5 Lasso Bayesian networks do not need accurate prior gene ordering

Interestingly, the performance of lasso-based networks did not depend strongly on the prior ordering, as shown in the comparisons between lasso-findr and lasso-random in Figure 3, Figure 4, and Supplementary Figure S7. Further inspections revealed a high overlap of top predictions by lasso-findr and lasso-random Bayesian networks, particularly among their true positives (Figure 5). This allows us to still recover genuine interactions even if the prior gene ordering is not fully accurate.

**Figure 5:**
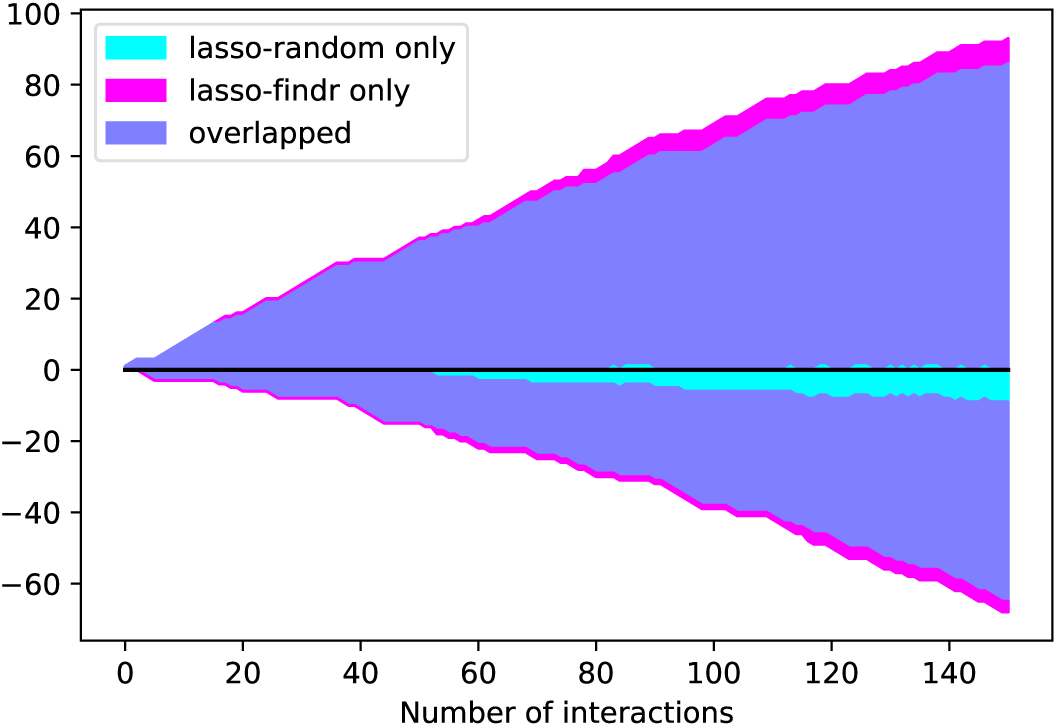
The numbers of overlap and unique interactions (*y* axis) predicted by lasso-findr and lasso-random Bayesian networks as functions of the number of significant interactions in each network (*x* axis), on DREAM dataset 1. Positive and negative directions in *y* correspond to true and false positive interactions according to the gold standard.

### 3.6 Lasso Bayesian networks mistake confounding as false positive interactions

We then tried to understand the differences between lasso and Findr based Bayesian networks, by comparing three types of gene relations in DREAM dataset 1, both among genes with a cis-eQTL in Figure 6A, and when also including genes without any cis-eQTL as only targets in Figure 6B. Both findr and lasso-findr showed good sensitivity for the genuine, direct interactions. However, when two otherwise independent genes are directly confounded by another gene, lasso tends to produce a false positive interaction, but not findr. As expected, to achieve optimal predictive performance, lasso regression cannot distinguish the confounding by a gene that is either unknown or ranked lower in the DAG.

**Figure 6:**
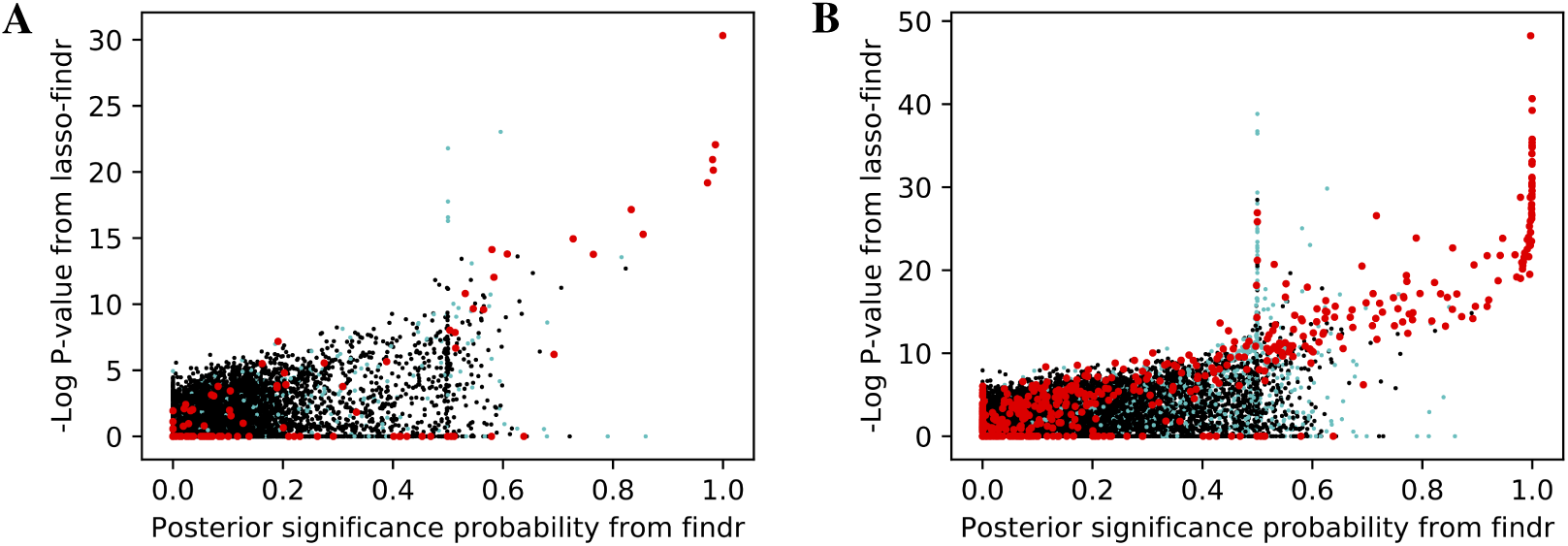
The significance score of findr (posterior probability; x-axis) and in lasso-findr (-log P-value; y-axis) for direct true interactions (red), directly confounded gene pairs (cyan), and other, unrelated gene pairs (black) on DREAM dataset 1; in **A**, only genes with cis-eQTLs are considered as regulator or target, whereas in **B** targets also include genes without cis-eQTLs. Higher scores indicate stronger significances for the gene pair tested.

### 3.7 Findr and lasso Bayesian network inference is highly efficient

The findr and lasso Bayesian networks required much less computation time compared to the bn-learn Bayesian networks, therefore allowing them to be applied on much larger datasets. To infer a Bayesian network of 230 genes from 100 samples in DREAM dataset 1, Findr required less than a second, lassopv around a minute, but bnlearn Bayesian networks took half an hour to half a day (Table 1). Moreover, since bnlearn only produces binary Bayesian networks, multiple recomputation is necessary to acquire the desired network sparsity.

**Table 1:** Timings for different Bayesian network inference methods/programs. Times for bnlearn methods depend on parameter settings (e.g. nominal FDR and AIC penalty), and take longer (approx. 8 times) with genotypes included. Times for bnlearn-hc include 10 random restarts.

| Dataset | Samples | Genes | findr | lassopv | bnlearn-hc | bnlearn-fi |
| --- | --- | --- | --- | --- | --- | --- |
| DREAM | 100 | 230 | <1sec | ≈1min | ≥10hr | ≥30min |
| Geuvadis | 360 | 3172 | <1min | ≈10hr | - | - |

### 3.8 Results on the Geuvadis dataset reaffirm conclusions from simulated data

To test whether the results from the DREAM data also hold for real data, we inferred findr and lasso-findr Bayesian networks from the Geuvadis data using both real and random causal priors (see Methods); conventional bnlearn-based network inference was attempted, but none of the restarts could complete within 1000 minutes.

Lasso-findr Bayesian networks were previously shown to provide ideal FDR control on this dataset [57], whereas findr Bayesian networks did not obtain a satisfying FDR control (Supplementary Figure S8). We believe this is due to the reconstruction of the node ordering, which interferes with the FDR control in pairwise causal inference. On the other hand, and again consistent with the DREAM data, findr Bayesian networks obtained superior results for the recovery of known transcriptional regulatory interactions inferred from ChIP-sequencing data (Figure 7A,B); neither method predicted TF targets inferred from siRNA silencing with high scores or accuracy better than random (Figure 7C).

**Figure 7:**
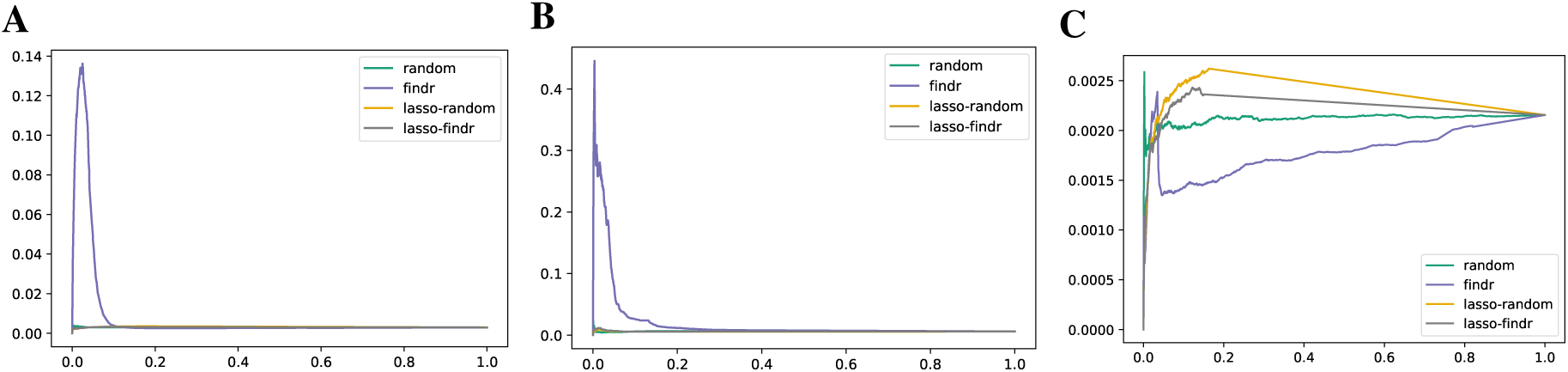
Precision-recall curves for Bayesian networks reconstructed from the Geuvadis dataset for three groundtruth networks: DNA-binding of 20 TFs in GM12878 (**A**), DNA-binding of 14 TFs in 5 ENCODE cell lines (**B**), and siRNA silencing of 6 TFs in GM12878 (**C**).

Comparisons on the predictive power yielded results similar with the DREAM datasets, where predictive scores were again hardly able to distinguish network directions.

## 4 Discussion

The inference of Bayesian gene regulatory networks for mapping the causal relationships between thousands of genes expressed in any given cell type or tissue is a challenging problem, due to the computational complexity of conventional hill-climbing, MCMC sampling or constraint-based methods. Here we have introduced an alternative method, which first reconstructs a topological ordering of genes, and then infers a sparse maximum-likelihood Bayesian network using variable selection of parents for every gene from its predecessors in the ordering. Our method is applicable when pairwise prior information is available or can be inferred from auxiliary data, such as genotype data. Our evaluation of the method using simulated genotype and gene expression data from the DREAM5 competition, and real data from human lymphoblastoid cell lines from the GEUVADIS consortium, revealed several lessons that we believe to be generalizable.

A major disadvantage of conventional score-based methods, irrespective of their computational cost, was their over-fitting of the expression profiles of a very small number of target genes. In high-dimensional settings where the number of genes far exceeds the number of samples, the expression profile of any one of them can be regressed perfectly (i.e. with zero residual error) on any linearly independent subset of variables, and this causes the log-likelihood to diverge. Even when the number of parents per gene was restricted to less than the number of samples, it remained the case that at any level of network sparsity, the divergence of the log-likelihood with decreasing residual variance of even a single gene resulted in score-based networks where most genes had either the maximum number of parents, or no parents at all. Restricting the maximum number of parents to an artificially small level can circumvent this problem, but will also distort the network topology, particularly by truncating the in-degree distribution, and therefore predict a biased gene regulatory network. Optimizing the total amount of variance explained, rather than log-likelihood, might overcome this problem. This, however, is not available yet in bnlearn.

Our method reconstructs a Bayesian network as a sparse subgraph from a maximum-weight DAG determined by pairwise causal relationships inferred using instrumental variable methods. We considered two variants of the method: one where the edge weights in the maximum-weight DAG were truncated directly to form a sparse DAG, and one where an additional L1-penalized lasso regression step was used to enforce sparsity. The lasso step was introduced for two reasons. First, pairwise relations do not distinguish between direct or indirect interactions and do not account for the possibility that a true relation may only explain a small proportion of target gene variation (e.g. when the target has multiple inputs). We hypothesized that adding a multi-variate lasso regression step could address these limitations. Second, truncating pairwise relations results in non-uniform false discovery rates for the retained interactions, due to each gene starting with a different number of candidate parents in the pairwise node ordering. As we showed in this paper and our previous work [57], a model selection p-value derived from lasso regression can control the FDR uniformly for each potential regulator of each target gene, resulting in an unbiased sparse DAG.

Despite these considerations, the ‘naive’ procedure of truncating the original pairwise causal probabilities resulted in Bayesian networks with better overlap with groundtruth networks of known transcriptional interactions, in both simulated and real data. We believe this is due to the lack of any instrumental variables in lasso regression, which makes it hard to dissociate true causal interactions from hidden confounding. Indeed, it is known that if there are multiple strongly correlated predictors, lasso regression will randomly select one of them [64], whereas in the present context it would be better to select the one that has the highest prior causal evidence. In a real biological system, findr networks and the use of instrumental variables may therefore be more robust than lasso regression, particularly in the presence of hidden confounders. We also note that the deviation from uniform FDR control for the naive truncation method was not huge and only affected genes with a very large number of candidate parents (Figure 2). Hence, at least in the datasets studied, adding a lasso step for better false discovery control did not overcome the limitations introduced by confounding interactions.

On the other hand, the lasso-random network used solely transcriptomic profiles, yet provided better performance than the conventional score-based and constrained-based networks, including those that used genotypic information. Together with its better false discovery control, this makes the lasso-random network an interesting method for high-dimensional Bayesian network inference with no or limited prior information.

In addition to comparing the inferred network structure against known ground-truths, we also compared the predictive performance of the various Bayesian networks. Although findr Bayesian networks again performed best, differences with lasso-based methods were modest. As is well known, using observational data alone, Bayesian networks are only defined upto Markov equivalence [21, 22], i.e. there is usually a large class of Bayesian networks with very different topology which all explain the data equally well. Hence it comes as no surprise that the prediction accuracy in edge directions has little impact on that in expression levels. This suggests that for the task of reconstructing gene networks, Bayesian network inference should be evaluated, and maybe also optimized, at the structural rather than inferential level. This also reinforces the importance of causal inference which, although challenging both statistically and computationally, demonstrated significant improvement of the global network structure even when it was restricted to pairwise causal tests.

Most of our results were derived for simulated data from the DREAM Challenges, but were qualitatively confirmed using data from human lymphoblastoid cell lines. This is because human groundtruth networks have strong limitations. They are normally reconstructed from heterogeneous, noisy, high-throughput data (e.g. ChIP-sequencing and/or knock-out experiments), and are both incomplete (many true interactions are not present) and imperfect (many detected physical interactions have no functional effect). In addition, statistical inference algorithms can hardly distinguish direct interactions from indirect ones, which operate through an unidentified third factor and should be regarded as “false positives”. As such, one has to be cautious not to over-interpret results, for instance on the relative performance of findr vs. lasso-findr Bayesian networks. Much more comprehensive and accurate ground-truth networks of direct causal interactions, preferably derived from a hierachy of interventions on a much wider variety of genes and functional classes (not only transcription factors), would be required for a conclusive analysis. Emerging large-scale perturbation compendia such as the expanded Connectivity Map, which has profiled knock-downs or over-expressions of more than 5,000 genes in a variable number of cell lines using a reduced representation transcriptome [65], hold great promise. However, the available cell lines are predominantly cancer lines, and the relevance of the profiled interactions for systems genetics studies of human complex traits and diseases, which are usually performed on primary human cell or tissue types, remains unknown.

Lastly, we note that our study has focused on ground-truth comparisons and predictive performances, but did not evaluate how well the second part of the log-likelihood, derived from the genotype data [cf. eq. (4)], was optimized. This score is never considered in the conventional score-based algorithms, and hence a comparison would not be fair. Moreover, optimising it is known to be an NP-hard problem. We used a common greedy heuristic optimization algorithm, but for this particular problem, this heuristic has no strong guaranteed error bound. We intend to revisit this problem, and investigate whether other graph-theoretical algorithms, perhaps tailored to specific characteristics of pairwise interactions inferred from systems genetics data, are able to improve on the greedy heuristic.

To conclude, Bayesian network inference using pairwise genetic node ordering is a highly efficient approach for reconstructing gene regulatory networks from high-dimensional systems genetics data, which outperforms conventional methods by restricting the super-exponential graph structure search space to acyclic graphs compatible with the causal inference results, and which is sufficiently flexible to integrate other types of pairwise prior data when they are available.

## Supplementary Information

### S1 Theoretical background and results

#### S1.1 Bayesian network primer

We collect here the minimal background on Bayesian networks necessary to make this paper selfcontained. For more details and proofs of the statements below, we refer to existing textbooks, for instance [21].

A Bayesian network for a set of continuous random variables *X*_1_, …, *X*_*n*_, represented by nodes 1, …, *n*, is defined by a DAG *𝒢* and a joint probability density function that decomposes as in eq. (1). We are interested in linear Gaussian networks, which can be defined alternatively by the set of structural equations

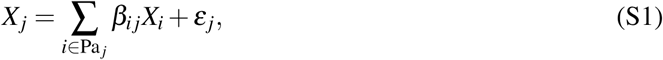

where Pa _*j*_ is the set of parent nodes for node *j* in *𝒢* and 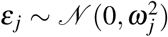 are mutually independent normally distributed variables. The matrix **B** = (*β*_*ij*_), with *β*_*ij*_ = 0 for *i* ∉ Pa_*j*_ can be regarded as a weighted adjacency matrix for *𝒢*. With this notation, the conditional distributions in eq. (1) satisfy

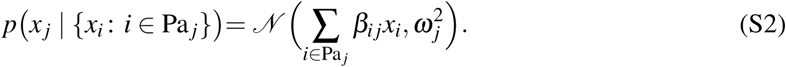

The values of **B** and 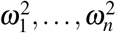 are the parameters of the Bayesian network which are to be determined along with the structure of *𝒢*. The conditional distributions (S2) result in the joint probability density function being multi-variate normal,

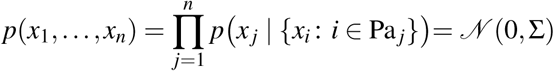

with inverse covariance matrix

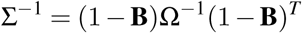

Where 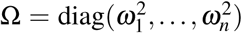. It follows that the gene expression-based term in the log-likelihood [eq. (6)] can be written as (up to an additive constant)

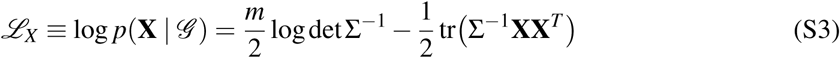

where as before **X** ∈ R^*n*×*m*^ is the data matrix for *n* genes in *m* independent samples. From these basic results, the following can be derived easily:

- For a given *𝒢*, *ℒ*_*X*_ can also be written as

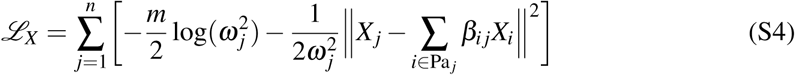

where *X*_*j*_ ∈ R^*m*^ is the expression data vector for gene *j*. It follows that the maximum-likelihood parameter values 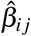 are the ordinary least-squares linear regression coefficients, 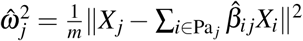 are the residual variances, and *ℒ*_*X*_ evaluated at these maximum-likelihood values is the log of the total unexplained variance, up to an additive constant

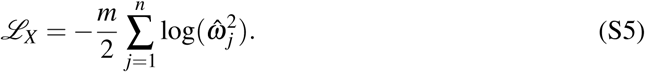
- Adding more explanatory variables always reduces the residual variance in linear regression. Hence 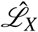 as a function of *𝒢* is maximized for fully connected DAGs with *n*(*n-* 1)*/*2 edges^1^. A fully connected DAG *𝒢* defines a “topological” node ordering *≺* by the relation

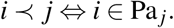

Equivalently, a node ordering defines a permutation *π* such that nodes are ordered as *π*_1_ *≺ π*_2_ *≺* …*≺ πn*. Permuting the rows and columns of **B** according to *π* turns **B** into a lower triangular matrix. Hence eq. (S5) can also be seen as a function on node orderings or permutations, and the maximum-likelihood values are then found by linearly regressing each node on its predecessors (i.e. parents) in the ordering [cf. eq. (S4)]. More precisely, we write

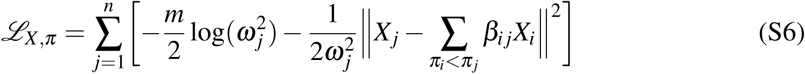
- Conversely, eq. (S3), and hence also eq. (S5), is easily seen to be invariant under any reordering of the nodes. Hence no edge directions can be inferred unambiguously from observational expression data without further constraints or information.

#### S1.2 Pairwise node ordering

To infer Bayesian gene networks, we first consider the log-likelihood score (6) without sparsity constraints,

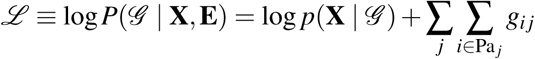

where it is implicitly understood that the maximum-likelihood parameters are used in *ℒ*_*X*_ = log *p*(**X** | *𝒢*). Because *ℒ*_*X*_ and 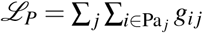 are both maximized for fully connected DAGs, and because the value of *ℒ*_*X*_ is the same for all fully connected DAGs, it follows that to maximize *ℒ*, we need to find the maximum-weight DAG which maximizes the pairwise score *ℒ*_*P*_. As stated in the main text, this is an NP-hard problem with no known polynomial approximation algorithms with a strong guaranteed error bound. The greedy algorithm we used is the standard heuristic for this type of problem [54].

#### S1.3 Sparsity constraints

Using fully connected DAGs leads to overfitting of the expression-based score *ℒ*_*X*_, particularly in the case where the number of genes *n* is greater than the number of samples *m*. The most popular methods for imposing sparsity in Bayesian networks are:

- **Bayesian or Akaike Information Criterion.** The BIC or AIC methods augment the likelihood function *ℒ*_*X*_ with a term proportional to the number of parameters in the model, i.e. the number of edges |*𝒢* | in *𝒢* (BIC = *-*|*𝒢* | log *m*, AIC = *-*|*𝒢* |).
- **L1-penalized lasso regression.** In this case, the likelihood *ℒ*_*X,π*_ [eq. (S6)] is augmented by a term 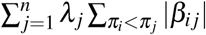, such that finding the maximum-likelihood parameters 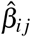 becomes equivalent to performing a series of independent lasso regressions, one for each node on its predecessors in the ordering *π*. The extra penalty term can be understood as coming from a double-exponential prior distribution on the parameters *β*_*ij*_.

An under-appreciated drawback of the BIC/AIC in high-dimensional settings is the fact that with a sufficient number of predictors it is possible to reduce 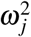 to zero for any gene, and hence make *ℒ*_*X*_ (S5) arbitrarily large. By concentrating all interactions on one or a few target genes, this can be achieved while still keeping the BIC/AIC small. Hence in high-dimensional settings, use of the BIC/AIC leads to highly skewed ‘all-or-nothing’ in-degree distributions, as shown in Figure 2C, unless the maximum allowed number of regulators for each gene is capped at an artificially small number.

Similar problems can occur if lasso regression is used with a fixed *λ* for all *j*, because the number of candidate regulators differs greatly among genes that come early or late in the ordering. In [38], a method was proposed where the value of *λ*_*j*_ increases with the order of *j*, but their scaling could not provide any guarantee for the probability of false positive errors for individual edges in the resultant sparse graph. We used the lassopv variable selection method [57] instead. In brief, for each gene *j* and for each candidate regulator *i* of *j* (i.e. predecessor of *j* in the ordering *π*):

- calculate the largest (most stringent) value of *λ*_*j*_ for which *i* would be selected as a parent of *j* (i.e. have non-zero lasso regression coefficient);
- calculate the probability (p-value) of a randomly generated predictor having the same or larger ‘critical’ *λ*_*j*_.

This results in a set of p-values *p*_*ij*_ for all pairs *π*_*i*_ < *π*_*j*_, which achieve optimal false discovery control, i.e.they can be transformed into q-values *q*_*i j*_ by standard FDR correction methods such that if we keep all *q*_*ij*_ ≤ *α*, the expected FDR is less than *α*. Moreover for sufficiently small thresholds *α*, there is a corresponding penalty parameter value *λ*_*j*_(*α*) such that the set of regulators with *p*_*i j*_ (or *q*_*i j*_) less than *α* is precisely the set of regulators with non-zero lasso regression coefficient [57]. Hence in our method we can use thresholding on the *p*_*i j*_ directly to obtain sparse Bayesian networks.

In addition to the lasso regression based method for inducing sparsity, we also considered a simple **thresholding on the pairwise prior information** to obtain a sparse DAG. In eq. (6), if we set

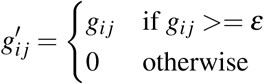

then edges with *g*_*ij*_ *< ε* are automatically excluded from the maximum-likelihood DAG, and the pairwise node ordering procedure will automatically result in a sparse DAG. This method does not provide any guarantee for the false positive control of individual edges in the (multi-variate) Bayesian network beyond what is provided by the pairwise causal inference test used.

#### S1.4 Summary of terminology

The following terminology is used repeatedly in this paper:

- **“Node ordering”:** a permutation of the nodes.
- **“Edge constraint”**: a set of ordered node pairs *C* = {(*i, j*)} in a DAG *𝒢*, that constrains the edges permitted in *𝒢* as ∀ *i* ∈ Pa _*j*_, (*i, j*) ∈*C*. Each DAG can be subject to more than one edge constraint.
- **“Topological node ordering”**: a node ordering *≺* to a DAG *𝒢*, that acts as an edge constraint *C = {*(*i, j*) | *i ≺ j}*.
- **“Fully connected DAG”**: a DAG *𝒢* in which no edge can be added. On a DAG with *n* nodes and with no edge constraint, it is fully connected if and only if it has *n*(*n -*1)*/*2 edges, because the addition of even a single edge is guaranteed to introduce a cycle, that is, *𝒢* would cease being a DAG. There is a one-to-one correspondence between node orderings and fully-connected DAGs.
- **“Maximum-weight DAG”**: a DAG that solves the maximum acyclic subgraph problem, identified by its parent sets

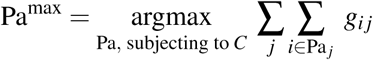

for some set of non-negative prior weights *g*_*ij*_. As discussed in the manuscript, there is no known algorithm for solving the maximum acyclic subgraph problem exactly. For simplicity, we also use the term “maximum-weight DAG” to refer to its heuristic, i.e. the local optima found by the greedy algorithm.

#### S1.5 Assessment of predictive power for Bayesian networks

The following 5-fold cross-validation algorithm was used to assess the predictive power of different Bayesian network inference methods.

##### Algorithm S1 Cross-validation of predictive power for Bayesian networks

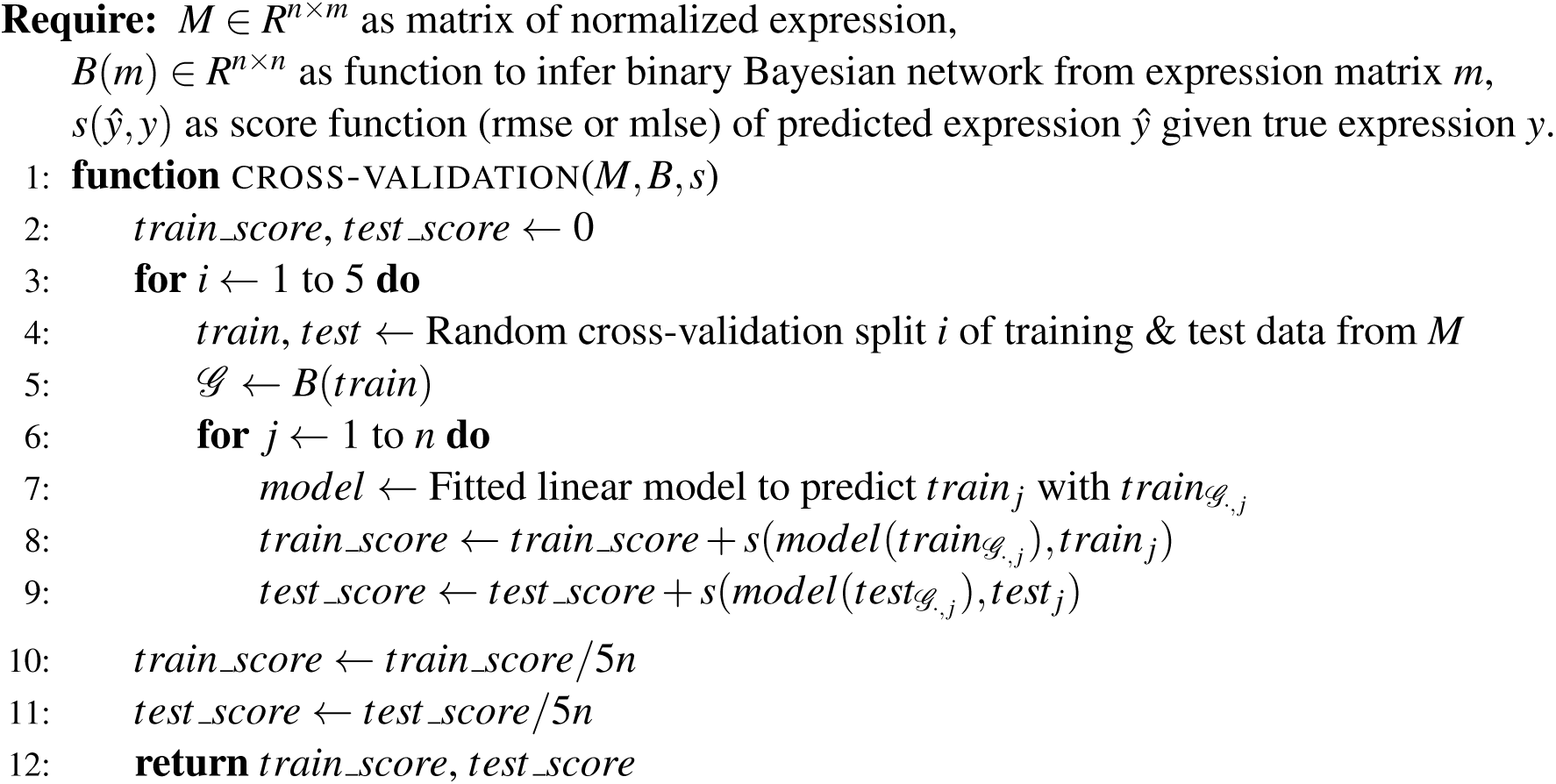

### S2 Supplementary figures and table

**Figure S1:**
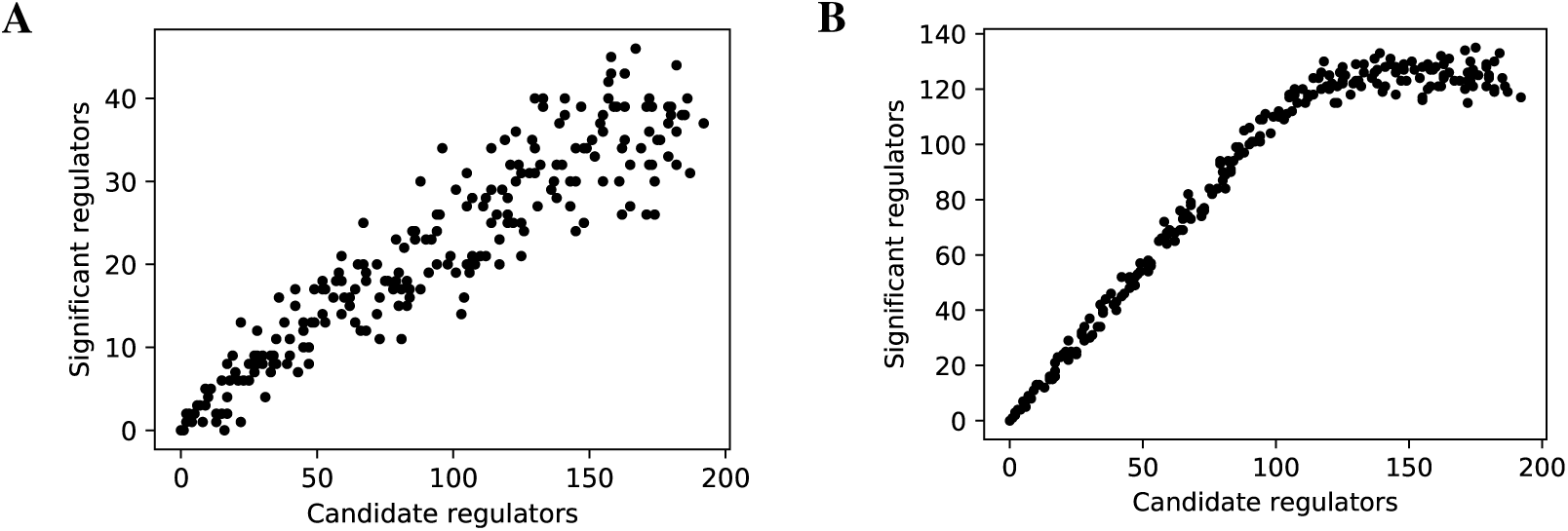
The linearity test of lasso-findr Bayesian networks at 5,000 (**A**) and 20,000 (**B**) significant interactions on DREAM dataset 1.

**Figure S2:**
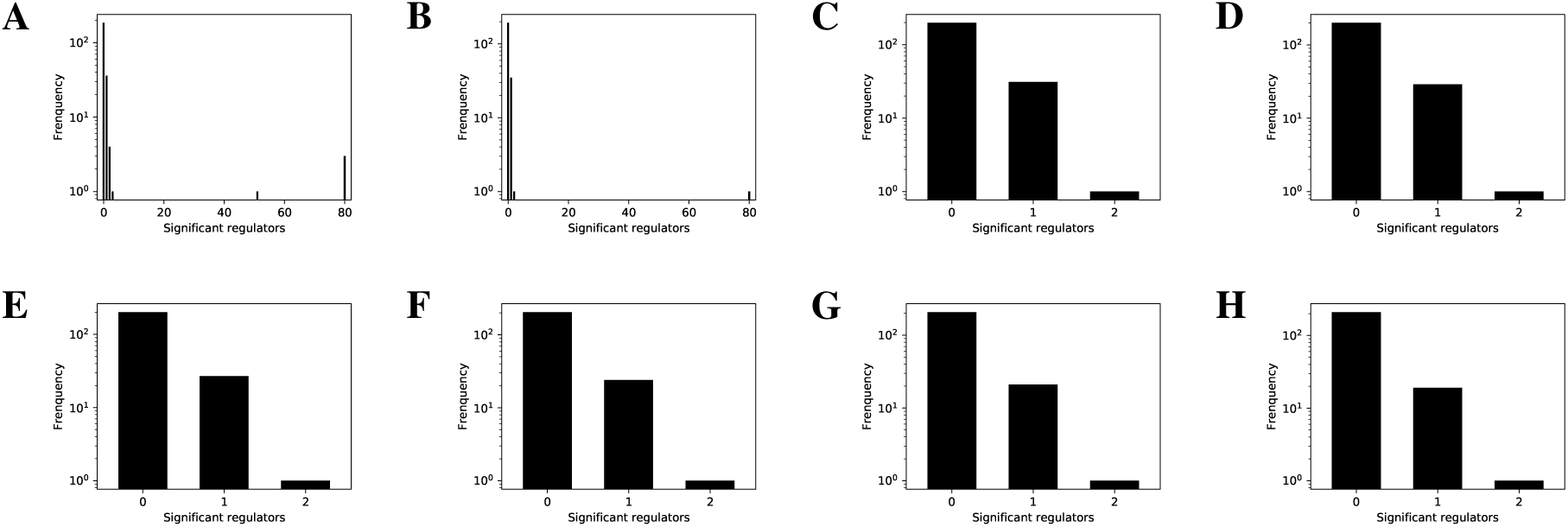
The histogram of significant regulator counts for each target gene in the bnlearn-hc Bayesian networks with AIC penalty 8.5 to 12 (**A** to **H**) and step 0.5 on DREAM dataset 1.

**Figure S3:**
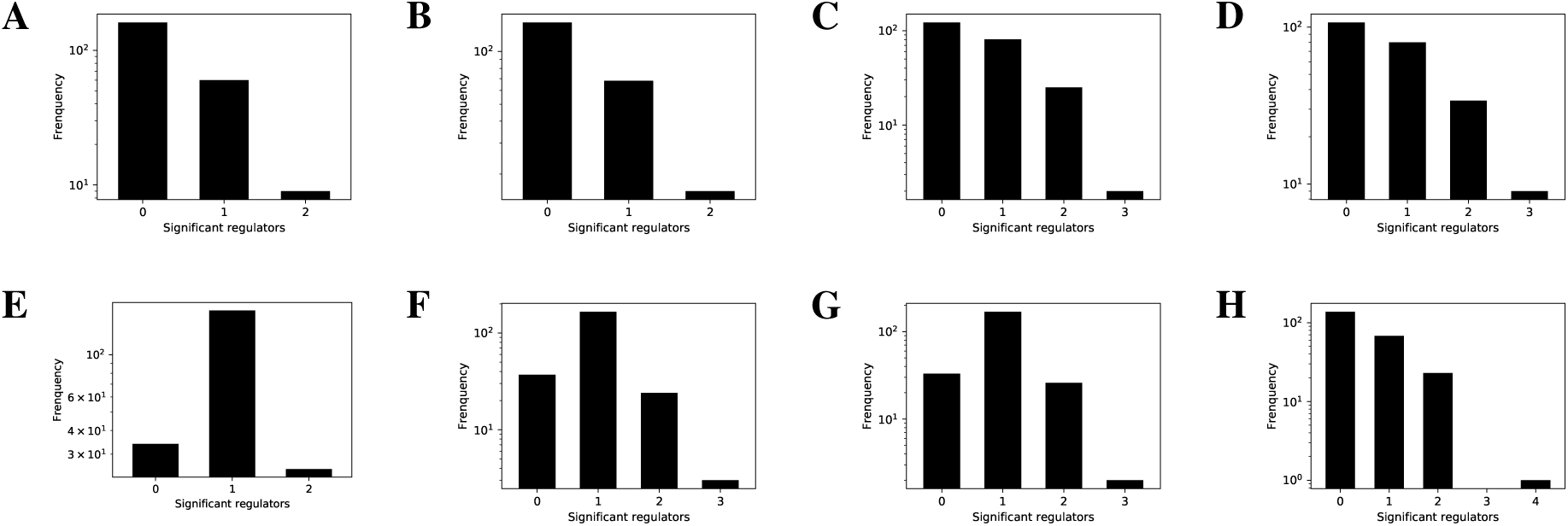
The histogram of significant regulator counts for each target gene in the bnlearn-fi Bayesian networks with nominal type I error rates 0.001, 0.002, 0.005, 0.01, 0.02, 0.03, 0.05, 0.2 (**A** to **H**) on DREAM dataset 1.

**Figure S4:**
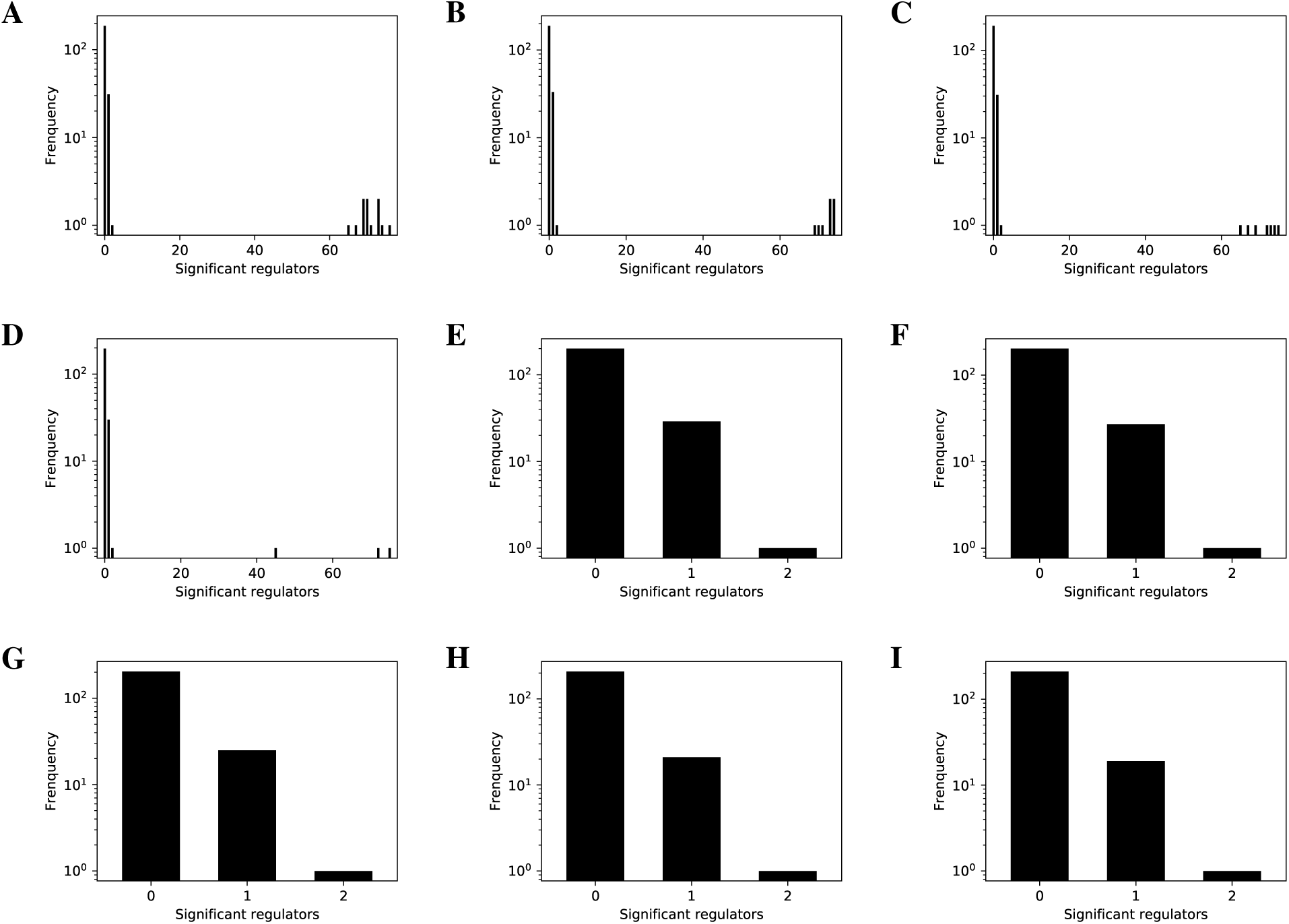
The histogram of significant regulator counts for each target gene in the bnlearn-hc-g Bayesian networks with AIC penalty 9.5 to 13 (**A** to **I**) and step 0.5 on DREAM dataset 1.

**Figure S5:**
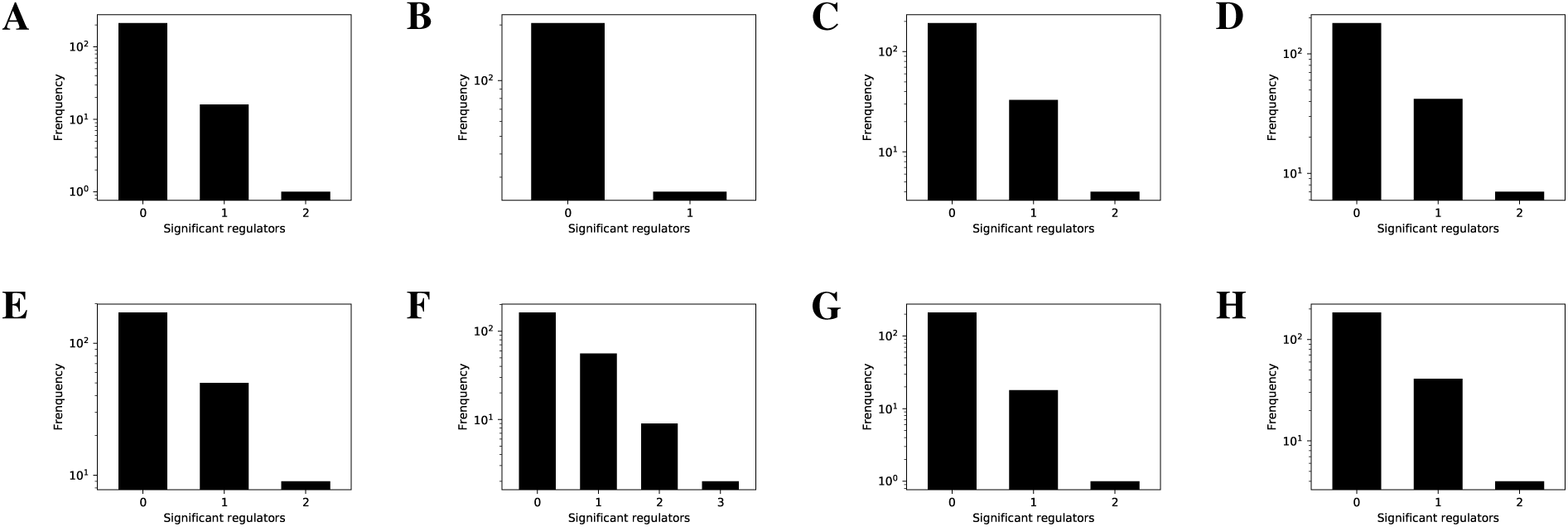
The histogram of significant regulator counts for each target gene in the bnlearn-fi-g Bayesian networks with nominal type I error rates 0.001, 0.002, 0.005, 0.01, 0.02, 0.03, 0.05, 0.2 (**A** to **H**) on DREAM dataset 1.

**Figure S6:**
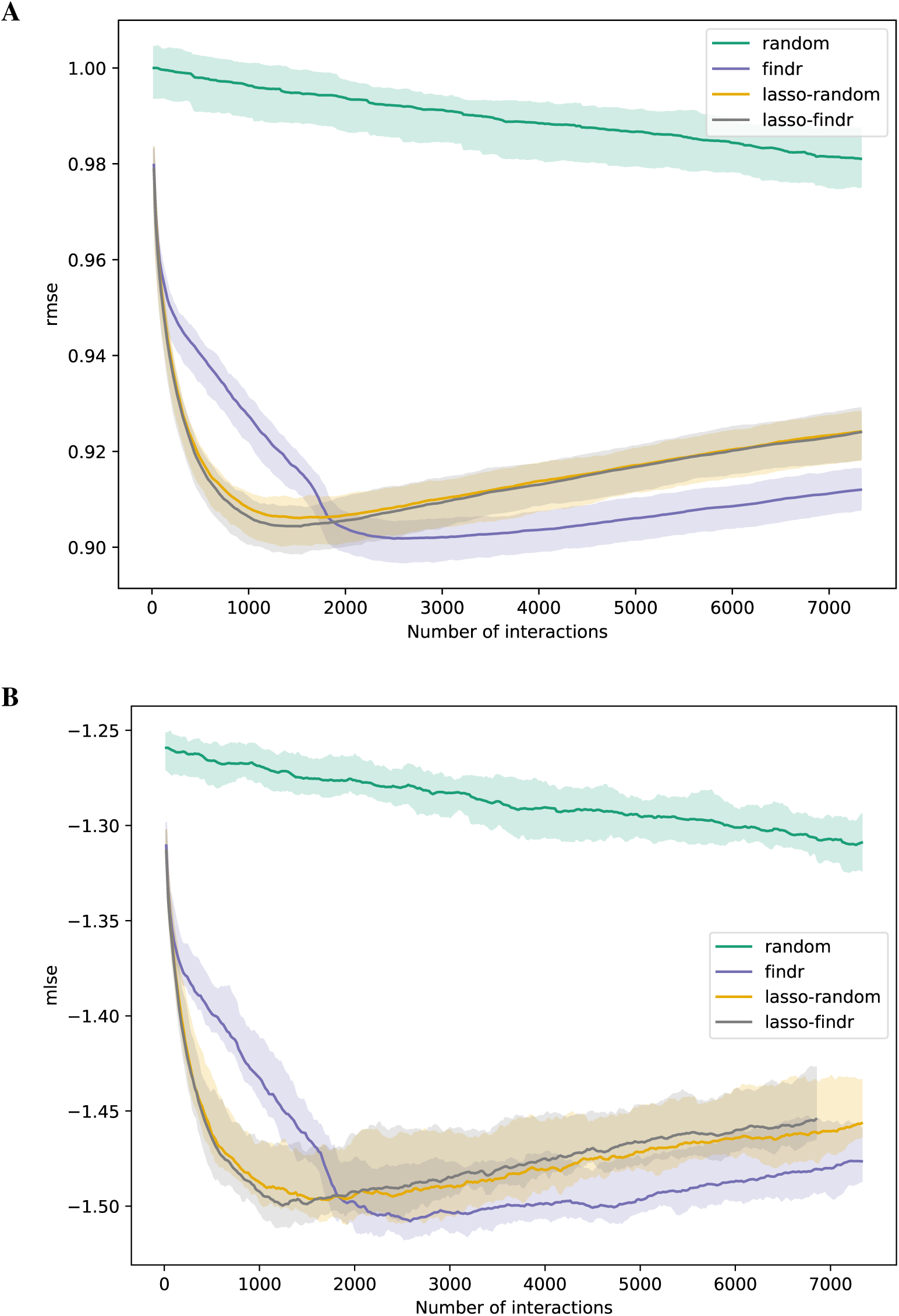
The root mean squared error (rmse, **A**) and mean log squared error (mlse, **B**) in training data are shown as functions of the numbers of predicted interactions in five-fold cross validations using linear regression models. Shades and lines indicate minimum/maximum values and means respectively. DREAM dataset 1 with 999 samples was used.

**Figure S7:**
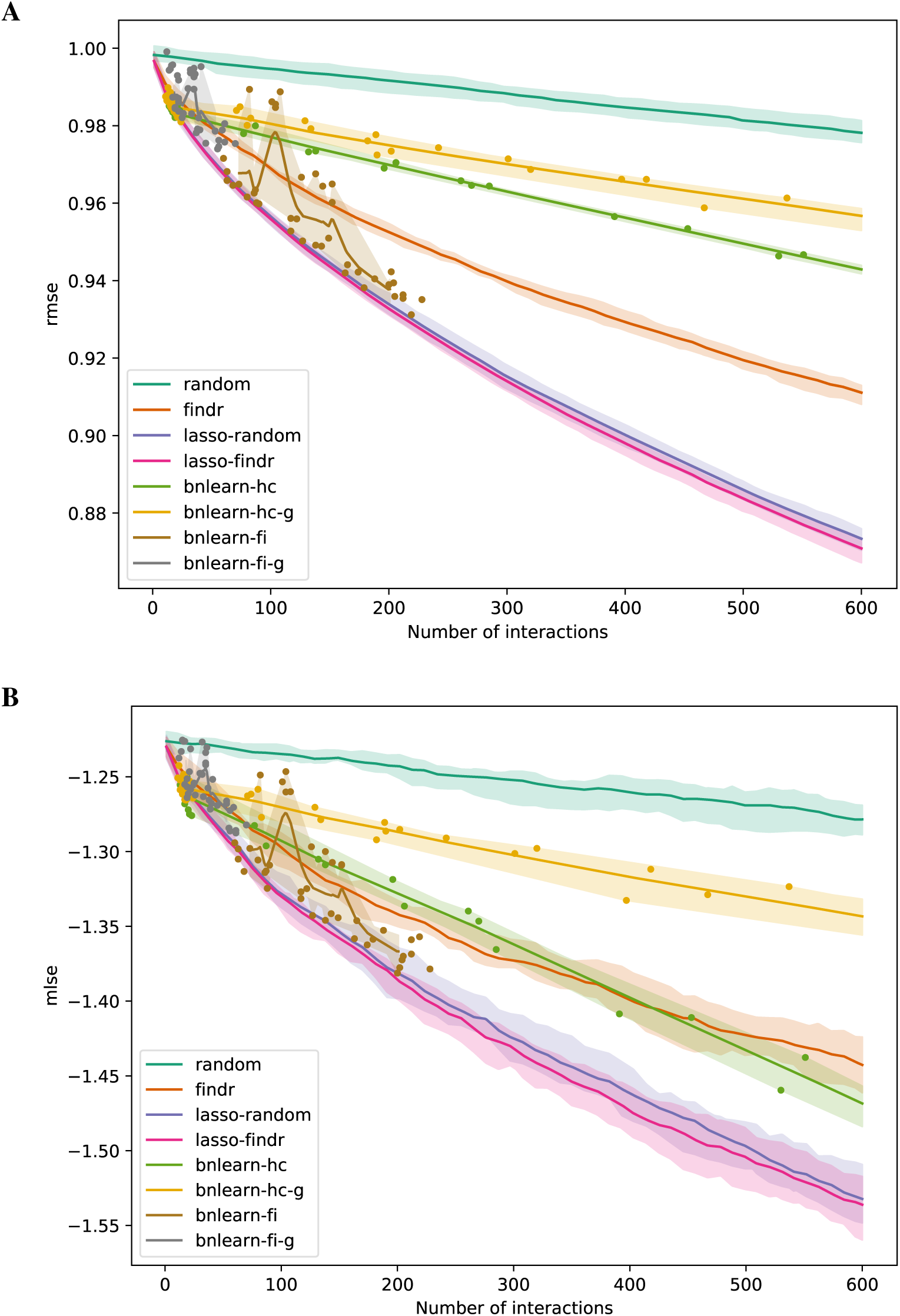
The root mean squared error (rmse, **A**) and mean log squared error (mlse, **B**) in training data are shown as functions of the numbers of predicted interactions in five-fold cross validations using linear regression models. Shades and lines indicate minimum/maximum values and means respectively. Root mean squared errors greater than 1 indicate over-fitting. DREAM dataset 1 with 100 samples was used.

**Figure S8:**
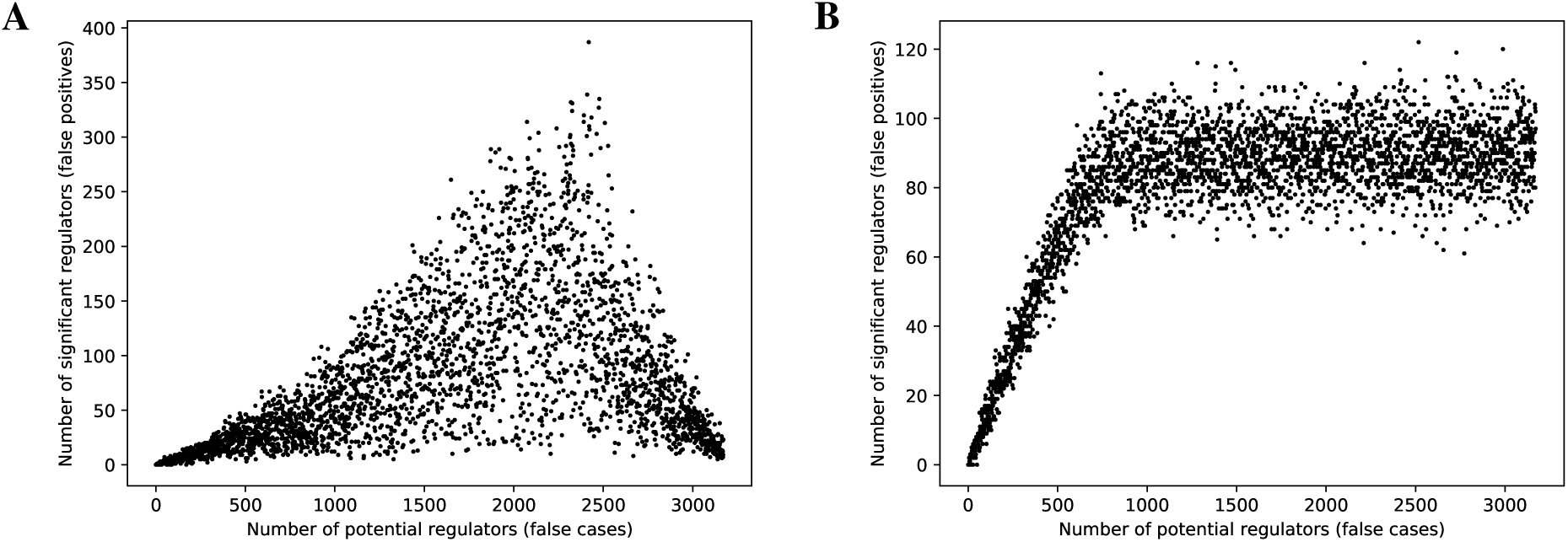
Conversion to Bayesian network from findr’s predictions breaks its false discovery control.

**Table S1:** Main characteristics of Bayesian network inference strategies considered in this paper.

|  | **Genetic node ordering** | **Score-based** | **Constraint-based** |
| --- | --- | --- | --- |
| **Algorithm goal** | Local optimum of log-likelihood | Local optimum of log-likelihood | Graph skeleton representing conditional independences |
| **Genotype data** | Causal inference between gene pairs | As extra parentless variables | As extra parentless variables |
| **Edge direction** | From causal inference step | Selected during hill-climbing | From resolution of skeleton constraints |
| **Search space restriction** | Only BNs compatible with node ordering induced by causal inference | Bounded in-degree | Bounded in-degree |
| **Sparsity constraint** | Variable selection with FDR control on node ordering | AIC | Nominal Type I error |

## Footnotes

1 We use the terminology “fully connected DAG” because *(i)* there exist DAGs with *n*(*n* ‒1)*/*2 edges, and *(ii)* any graph with more than this number of edges contains at least one cycle, that is, is not a DAG.

